# Prevalence and risk factors of ovine and caprine fasciolosis in Ethiopia: a systematic review and meta-analysis

**DOI:** 10.1101/2025.04.23.650175

**Authors:** Simachew Getaneh Endalamew, Alebachew Tilahun Wassie, Andnet Yirga Assefa, Yihenew Getahun Ambaw, Solomon Mekuriaw Ayalew, Solomon Keflie Assefa

## Abstract

**Background:** Fasciolosis is a parasitic disease caused by liver flukes of the genus *Fasciola*, predominantly *Fasciola hepatica* and *Fasciola gigantica.* This zoonotic disease significantly impacts both livestock and human populations, particularly in areas with extensive agriculture and poor sanitation. Ethiopia, one of the Africa’s leading ovine and caprine producers, is highly affected by fasciolosis. However, despite its economic and public health importance, there is a lack of comprehensive up-to-date evidence on prevalence and risk factors of small ruminant fasciolosis. Therefore, this systematic review and meta-analysis aims to explore the pooled prevalence of fasciolosis among small ruminants (ovine and caprine) in Ethiopia.

**Methods:** This systematic review and meta-analysis were conducted based on the Preferred Reporting Items for Systematic Reviews and Meta-Analyses (PRISMA) guidelines. A comprehensive systematic review was performed across five electronic databases (Google Scholar, Embase, PubMed, Web of Science, and ScienceDirect), with all database searches and registers inquiries finalized on November 26, 2024. A random-effect model was used to determine the pooled prevalence of fasciolosis in ovine and caprine. Heterogeneity was assessed, and the source of variation was analyzed using subgroup and sensitivity analysis. Publication bias assessment and meta-regression analysis were also performed to ensure the robustness of the review. Funnel plots and Egger’s asymmetry tests were used to investigate publication bias.

**Results:** Overall, 33 studies were included in the meta-analysis, and the pooled prevalence of fasciolosis in ovine and caprine was 32.25% (95% CI: 25.97–38.86%). This study revealed substantial between-study heterogeneity (inconsistency index (I^2^)) = 97.3%, p < 0.0001). Among the variables analyzed for heterogeneity, species, publication years, season of data collection, and regions of the study were the most significant predictors of heterogeneity The sub-group analysis showed that the prevalence of fasciolosis among ovine and caprine was 37.18% (95% CI; 31.06–43.51%) and 12.76% (95% CI; 4.06–25.19%), respectively. According to the region-based subgroup meta-analysis, studies taken from Amhara region had the highest prevalence of fasciolosis among small ruminants (43.99% (95% CI: 31.83–56.52%)).

**Conclusion:** This systematic review and meta-analysis emphasize fasciolosis as a pervasive threat to Ethiopian small ruminants. Policymakers, healthcare providers, and communities should collaborate to integrate robust prevention mechanisms for the disease, establishing standardized protocols for *Fasciola* monitoring, reporting, and mitigation.

## Introduction

Small ruminant fasciolosis, which predominantly affects sheep and goats (Ovine and Caprine), is endemic to Ethiopia [1]. The life cycle of Fasciola depends on freshwater Lymnaea snails, which serve as intermediate hosts and thrive in riparian ecosystems such as riverbanks, irrigation channels, and marshy grazing lands [2]. Beyond small ruminants, fasciolosis results in substantial economic losses in cattle production and poses a zoonotic risk to humans via contaminated water or vegetables [3]. In Ethiopia, the prevalence of the disease is exacerbated by climatic factors (e.g., seasonal flooding), traditional pastoral practices, and limited veterinary intervention, creating a persistent health challenge [4].

Fasciolosis, a neglected parasitic infection caused by liver flukes (*Fasciola hepatica* and *Fasciola gigantica*), remains a critical global health and agricultural challenge [5, 6]. This zoonotic disease disproportionately affects communities in regions with poor sanitation and intensive agricultural practices, where transmission occurs through ingestion of water or food contaminated with metacercariae (the encysted form of the parasite) [7]. Recognized by the World Health Organization (WHO) as a neglected tropical disease, fasciolosis imposes severe socioeconomic burdens, particularly in low-resource settings where human and animal health systems are underdeveloped [8].

The life cycle of *Fasciola* species relies on *Mollusks or snails* of the family Lymnaeidae as intermediate hosts. Within these hosts, larval stages undergo multiple developmental phases and asexual replication before reaching maturity in definitive mammalian hosts, such as livestock and humans, where sexual reproduction occurs [9]. Transmission to definitive hosts occurs through ingestion of contaminated water, vegetation, or food containing metacercariae (the infective larval stage). Individuals with frequent exposure to infected livestock or contaminated environments face elevated risks of zoonotic transmission [7].

In livestock, chronic fasciolosis reduces productivity; causing weight loss, decreased milk yield, infertility, and liver condemnation, resulting in annual losses exceeding **$3.2 billion globally** [10–12]. Over **550 million ruminants,** including cattle, sheep, and goats, are affected worldwide, undermining food security and livelihoods [13, 14]. Human infections, historically overlooked, are increasingly reported in endemic regions, with complications such as hepatobiliary damage and chronic malnutrition exacerbating poverty cycles [15].

The habitats of the intermediate host (Mollusks) are climate-sensitive. Elevated transmission risks correlate with high rainfall, humidity, and moderate temperatures [16]. In Ethiopia, *F. hepatica* predominates in high-altitude areas (>1800m), while *F. gigantica* thrives in lowland regions (<1200m), though co-infections occur in ecologically overlapping zones [4] [17] [18]. Goats exhibit lower acute infection rates due to browsing behavior, but chronic infections in all ruminants silently diminish productivity, masking the disease’s economic toll [19].

Ethiopia stands as a leading producer of sheep and goats (ovine and caprine) in Africa, yet the country faces a critical gap in consolidated, nationwide data on small ruminant fasciolosis. While numerous localized studies have explored this parasitic infection over the past decade, their fragmented scope and geographic limitations hinder a unified understanding of its prevalence and risk factors. To address this, a systematic review and meta-analysis of research spanning 2015–2024 was conducted, synthesizing existing evidence to estimate a national pooled prevalence estimate and identify key epidemiological predictors.

This analysis provides actionable insights for veterinary health professionals, policymakers, and researchers, enabling evidence-driven strategies to optimize diagnostics, refine treatment protocols, and prioritize region-specific interventions. Furthermore, these findings contribute to global frameworks aimed at reducing the burden of zoonotic diseases, emphasizing the intersection of animal health, agricultural sustainability, and public health resilience.

## 2. Methods and analysis

### 2.1. Development of the review method

This systematic review and meta-analysis employed the Condition, Context, and Population (CoCoPop) framework [20] to structure its design. Adhering to the Preferred Reporting Items for Systematic Reviews and Meta-Analyses (PRISMA) Protocols 2020 guidelines [21], the methodology ensured rigor in protocol development, execution, and reporting. Prior to data collection, the study protocol was prospectively registered with PROSPERO (International Prospective Register of Systematic Reviews) on 21 February 2024 (Registration ID: CRD42024576654), with a focus on fasciolosis in sheep and goats. The analysis specifically evaluated the pooled prevalence of fasciolosis in these species as the primary outcome.

### 2.2. Inclusion and exclusion criteria

All cross-sectional studies published in peer reviewed journals that were conducted in Ethiopia were eligible for inclusion in this study. To ensure the incorporation of current and pertinent evidence on the prevalence of fasciolosis in small ruminants, only studies conducted after 2015 were included. Different study formats that reported the prevalence of fasciolosis in small ruminants, including peer-reviewed journal articles, Master’s theses, and dissertations published in English were considered.

Research articles were excluded for one of the following reasons: (a) reported the knowledge, attitudes, and practices of small ruminant fasciolosis (qualitative studies), (b) articles with insufficient information or records with missing outcomes of interest or poor-quality articles, (c) personal opinions, correspondence, letters to the editor, proceedings, and reviews. In addition, to enhance the study’s precision, articles with small sample sizes (less than 100) were excluded.

### 2.3. Searching strategy

Data was retrieved from five major academic databases, including Google Scholar (scholar.google.com), Embase (embase.com), PubMed (pubmed.ncbi.nlm.nih.gov), Web of Science (thomsonreuters.com), and ScienceDirect (sciencedirect.com). The final literature search was finalized on November 26, 2024.

The MeSH terms and keywords combined with Boolean operators were used to retrieve all relevant articles from the registers and databases. The search included keywords related to the population (“small ruminants”, “ovine”, “caprine”, “sheep”, “goat”), condition (“fasciolosis”, “*F. gigantica* “, “*F. hepatica* “) and context (e.g., “Ethiopia”). Boolean operators such as “AND” and “OR” were also applied to combine terms effectively. The search strategy, initially developed for PubMed, was adapted for other databases and included the following framework: ((((((((((((Ethiopia[Text Word]) OR (Ethiopia[MeSH Terms])) AND (sheep[MeSH Terms])) OR (sheep[Text Word])) OR (goat[MeSH Terms])) OR (goat[Text Word])) OR (small ruminants[Text Word])) AND (fasciolosis[MeSH Terms])) OR (*F. gigantica*[Text Word])) OR (*F. hepatica*[Text Word])) AND (prevalence[Text Word])) OR (seroprevalence[Text Word])). Retrieved records were organized using EndNote reference management software, where duplicates were eliminated through manual verification.

### 2.4. Evaluation of the quality of studies and risk of bias assessment

The quality of the included studies was assessed using Joanna Briggs Institute’s critical appraisal tool, designed to evaluate prevalence studies [20]. The tool has nine items measured with three options: “yes,” “no,” and “unclear”. The assessment of included studies was conducted by two independent teams: Team A (ATW, SGE, SMA) and Team B (SKA, YHA, AYA). These teams utilized a three-tier evaluation system (“yes,” “no,” “unclear”) to appraise article quality. Disagreements between evaluators were resolved through collaborative discussions and consultations with a panel of expert researchers. In the scoring protocol, a value of 1 was assigned for each “yes” response, while “no” or “unclear” answers (including instances where information was not reported or deemed irrelevant) received 0 points. Cumulative scores for each article were calculated on a scale of 0 to 9. Based on these totals, studies were classified into three quality orders: high (7–9 points), moderate (4–6 points), or low (0–3 points). For this systematic review and meta-analysis, only articles ranked as high or moderate quality were retained for final analysis.

### 2.5. Extraction of data from eligible papers

The initial search results passed relevance screening, with only studies meeting predefined inclusion criteria proceeding to the analysis phase. A standardized extraction template was utilized for data collection, implemented independently by two research teams: Team A (ATW, SGE, SMA) and Team B (SKA, YHA, AYA). Discrepancies in extracted information were resolved through consensus-based discussions between the groups. The template captured key study parameters including author name, publication year, geographic location, diagnostic method, sample size, climatic conditions, season of data collection, prevalence rates, and elevation data. All extracted information was systematically organized and cross-verified using Microsoft Excel (version 16.54) to ensure accuracy and consistency throughout the review process.

### 2.6. Data synthesis and Statistical analysis

Data extraction focused on compiling proportions, standard errors, and 95% confidence intervals (CIs) from studies reporting relevant findings. For studies lacking precomputed 95% CIs, these intervals were derived using validated statistical formulas [22]:

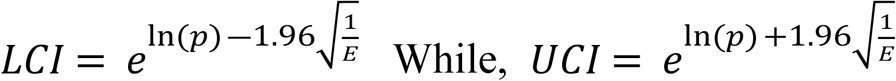

Here, *p* represents the proportion of fasciolosis cases, *E* denotes the total count of identified *Fasciola* specimens, and LCI/UCI define the interval bounds. This approach ensured methodological consistency and accuracy in estimating uncertainty across heterogeneous datasets.

Statistical analyses were conducted using R Studio (v4.4.2) with the “Meta” package (v8.0-1). Model selection (fixed vs. random effects) was guided by heterogeneity assessment, where a random effects framework was adopted due to significant between-study variability (I² = 97.3%). To optimize data normalization, five transformation methods were evaluated: untransformed prevalence rates (PR), arcsine (PAS), double-arcsine (PFT), logarithmic (PLN), and logit (PLOGIT). Model fitn1ess was determined using Shapiro-Wilk normality criteria, prioritizing transformations with W-statistics approaching 1 and non-significant p-values (>0.05). The double-arcsine method (PFT) demonstrated optimal alignment with normality assumptions (W = 0.95291, p = 0.1618), confirming its suitability for the final meta-analytic model.

A random-effects meta-analysis was conducted to account for variability across studies, with variance estimation performed through the DerSimonian and Laird approach [23]. PFT-transformed prevalence rates of small ruminant fasciolosis and associated standard errors were calculated for each study using the inverse variance method, facilitating pooled estimates [22, 24]. To enhance uncertainty quantification, confidence intervals were adjusted via the Hartung-Knapp-Sidik-Jonkman (HKSJ) procedure [25, 26]. The extent of between-study heterogeneity was further assessed by computing confidence intervals for τ² (tau-squared) and τ (tau) using the Jackson method, which quantified both the magnitude and precision of variability among included studies [27].

The distribution of fasciolosis prevalence among small ruminants was graphically synthesized using a forest plot, displaying individual study estimates with 95% confidence intervals (CI). Between-study heterogeneity, attributable to methodological diversity, was quantified using the Cochrane Q-test [28], I² statistic (proportion of total variability due to heterogeneity rather than sampling error) [29], and prediction intervals to estimate the potential range of true effects [30]. Subgroup analysis were conducted based on sample size, geographic region, animal species studied, season, publication year, and altitude to explore sources of variability. Sensitivity analyses assessed the reliability of combined results by systematically removing each study one at a time. To identify factors influencing variability, season, publication year, sample size, and region were analyzed using both univariable and multivariable meta-regression models within a random-effects structure. Baujat plots were additionally employed to identify studies that had a significant impact on overall variability. Publication bias was assessed by checking the symmetry of funnel plots and performing Egger’s regression test. A non-significant outcome in Egger’s test (*p* > 0.05) suggested no evidence of significant publication bias or small-study effects.

## 3. Results

### 3.1. Study selection and identification

Of the 2, 116 studies initially screened, 245 were removed due to duplication. Another 760 studies were excluded based on their titles and abstracts, as they were considered irrelevant to the review. Additionally, 28 studies were excluded for reasons such as not reporting the full outcome/reviews, poor quality, and inadequate sample size. Finally, 33 studies were considered suitable for inclusion in the qualitative and quantitative syntheses, as depicted in the PRISMA flow diagram (Figure 1).

**Figure 1.**
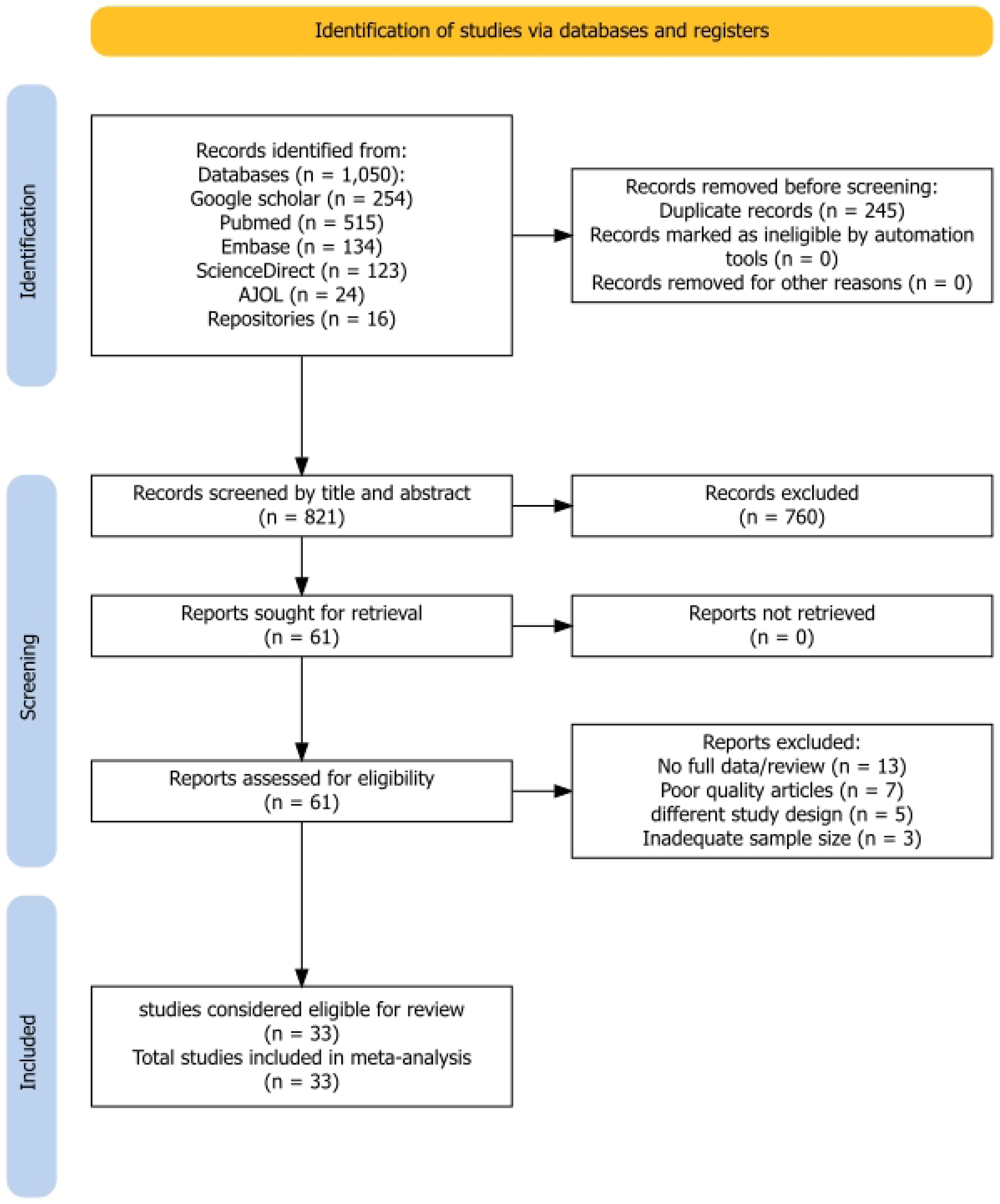
PRISMA flow diagram for study selection (identification, screening, eligibility assessment, and inclusion of studies) in the systematic review and meta-analysis

### 3.2. Description of included studies

This systematic analysis incorporated data from 33 studies conducted between 2015 and 2024 within field, clinical and abattoir settings, comprising a total sample of 9,578 small ruminants. Among these, 3,419 individuals (32.25%) exhibited confirmed fasciolosis disease. Between 2015 and 2024, the Oromia and Amhara region accounted for the majority of studies documenting small ruminant fasciolosis in Ethiopia. Most of the studies were conducted on ovine species. The sample sizes varied from 121 to 576 small ruminants. The apparent prevalence of the disease ranged from 5.80% to 70.2% in the study areas (Table 1).

**Table 1.** Characteristics of the included studies and prevalence of fasciolosis among small ruminants in Ethiopia, 2024 (n=33)

| Authors with publication years | Species | No. of Examined | No. of Positive | Prevalence with 95% CI | Diagnosis method used |
| --- | --- | --- | --- | --- | --- |
| Abaya, S. W., et al. (2023a) [31] | Ovine | 195 | 85 | 43.58 (35.24; 53.91) | Immunological test |
| Abaya, S. W., et al. (2023b) [31] | Ovine | 141 | 13 | 9.21 (5.35; 15.87) | Immunological test |
| Abdurahaman, M., et al. (2019) [32] | Ovine | 285 | 150 | 52.63 (44.84; 61.76) | Coproscopy |
| Alemu, B. and D. Chala (2019) [33] | Ovine | 384 | 151 | 39.32 (33.53; 46.13) | Coproscopy |
| Amsalu, T. (2017)[34] | Ovine | 576 | 61 | 10.59 (8.23; 13.62) | Coproscopy and Postmortem |
| Asrede, T. and A. Shifaw (2015) [35] | Ovine | 390 | 198 | 50.76 (44.16; 58.36) | Coproscopy |
| Ayele, Y. et al. (2018)[36] | Ovine | 280 | 170 | 60.71 (52.24; 70.57) | Coproscopy and Postmortem |
| Bayu, A. and S. Derso (2015) [37] | Ovine | 384 | 170 | 44.27 (38.09; 51.45) | Coproscopy |
| Bedada, H. and F. G. W. Negash (2017a) [38] | Ovine | 129 | 33 | 25.58 (18.18; 35.99) | Coproscopy |
| Bedada, H. and F. G. W. Negash (2017b) [38] | Caprine | 166 | 63 | 37.95 (29.65; 48.59) | Coproscopy |
| Belete, K. (2017) [39] | Ovine | 383 | 158 | 41.25 (35.29; 48.21) | Coproscopy |
| Berhanu, M., et al. (2016) [40] | Ovine | 384 | 189 | 49.21 (42.68; 56.77) | Coproscopy |
| Bimirew, F. L. and T. M. Cherinnat (2020) [41] | Ovine | 390 | 177 | 45.38 (39.16; 52.59) | Coproscopy |
| Birhanu, A., et al. (2015a) [42] | Ovine | 205 | 53 | 25.85 (19.76; 33.85) | Coproscopy |
| Birhanu, A., et al. (2015b) [42] | Caprine | 179 | 19 | 10.61 (6.78; 16.65) | Coproscopy |
| Chala, D. and B. Alemu (2019) [43] | Ovine | 384 | 151 | 39.32(33.53; 46.13) | Coproscopy and Postmortem |
| Dabasa, G., et al. (2017) [44] | Ovine | 384 | 88 | 22.91(18.6; 28.25) | Coproscopy |
| Demissie, T., et al. (2021a) [45] | Ovine | 168 | 28 | 16.66(11.51; 24.13) | Coproscopy and Postmortem |
| Demissie, T., et al. (2021b) [45] | Caprine | 216 | 26 | 12.03(8.19; 17.68) | Coproscopy and Postmortem |
| Destaw, K., et al. (2017) [46] | Ovine | 384 | 139 | 36.19(30.66; 42.75) | Coproscopy |
| Getabalew, M., et al (2019) [47] | Ovine | 121 | 85 | 70.24(56.8; 86.89) | Coproscopy |
| Ibrahim, A., et al. (2017) [48] | Ovine | 384 | 164 | 42.7(36.65; 49.78) | Coproscopy |
| Jarso D, e. a. (2016)[49] | Ovine | 475 | 229 | 48.21(42.35; 54.88) | Coproscopy and Postmortem |
| Kassye, D., et al. (2017a) [50] | Ovine | 202 | 75 | 37.12(29.61; 46.56) | Postmortem |
| Kassye, D., et al. (2017b) [50] | Caprine | 194 | 13 | 6.7(3.9; 11.55) | Postmortem |
| Kebede, S. and O. Wakgari (2016) [51] | Ovine | 384 | 214 | 55.72(48.75; 63.72) | Coproscopy |
| Megersa, B., et al. (2024a) [52] | Ovine | 220 | 54 | 24.54(18.8; 32.05) | Coproscopy |
| Megersa, B., et al. (2024b) [52] | Caprine | 112 | 12 | 10.71(6.08; 18.87) | Coproscopy |
| Melkamu, S. and M. Asrat (2015) [53] | Ovine | 384 | 137 | 35.67(30.17; 42.18) | Coproscopy |
| Mengistu, B. M., et al. (2019a) [54] | Ovine | 245 | 59 | 24.08(18.66; 31.09) | Coproscopy |
| Mengistu, B. M., et al. (2019b) [54] | Caprine | 139 | 8 | 5.75(2.88; 11.51) | Coproscopy |
| Regasa, A., et al. (2022) [55] | Ovine | 327 | 140 | 42.81(36.27; 50.53) | Coproscopy |
| Regea, G. and G. Getachew (2021) [56] | Ovine | 384 | 126 | 32.81(27.56; 39.07) | Coproscopy |
| Zekarias, T. and T. Bassa (2019) [57] | Ovine | 384 | 69 | 17.96(14.19; 22.76) | Coproscopy |

### 3.3. The pooled prevalence and prediction interval estimate of Fasciolosis in Ethiopia

This meta-analysis identified heterogeneity between the included studies (I^2^ = 97.3% (95% CI: [96.7%; 97.7%], p < 0.001), and *τ*^2^ = 0.0307 (95% CI: [0.0211; 0.0597], p < 0.001)). As a result, we used a random effects model to estimate the pooled prevalence of Fasciolosis. Based on the random-effects model, the pooled effect size of Fasciolosis among ovine and caprine was 0.3225 (95% CI: [0.2597; 0.3886] (Figure 2).

**Figure 2:**
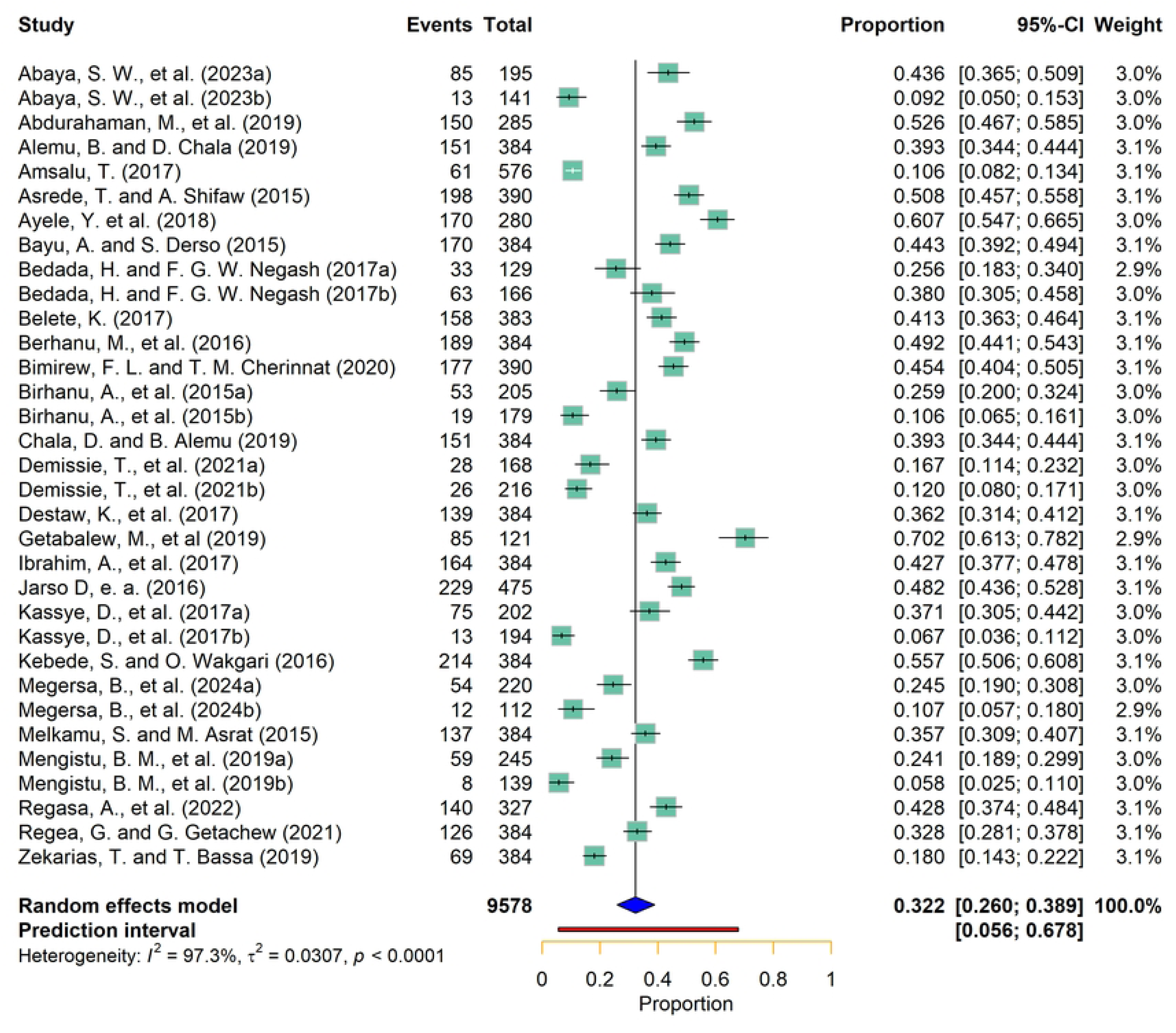
Forest plot for pooled prevalence of Fasciolosis in Ethiopia

A prediction interval estimates the range within which the results a future study, drawn from the same population, are expected to fall. This interval quantifies the expected uncertainty in the combined effect estimate when incorporating additional research into the meta-analysis [58, 59]. Based on the findings of this systematic review and meta-analysis, the predicted prevalence of fasciolosis among small ruminants in Ethiopia is expected to fall within the range of 5.6% to 67.8%. This suggests that future studies will likely report results within these intervals.

### 3.4. Handling heterogeneity

The meta-analysis indicated that between-study variability was high (Q = 1166.87, df = 32, p < 0.0001), indicating statistically significant differences in effect sizes across the included studies. The variance estimated was *τ*^2^ = 0.0307 (95% CI: [0.0211; 0.0597], with a *I*^2^ value of (I^2^ = 97.3% (95% CI: [96.7%; 97.7%]. Sensitivity analysis, subgroup analysis, and meta-regression analysis, were employed to address the substantial heterogeneity observed in the pooled estimate derived from the random-effects model. These methods help to explore potential sources of heterogeneity and provide insights into the robustness of the findings.

#### 3.4.1. Subgroup analysis

Subgroup analysis was conducted for the diagnostic methods used (Coproscopy, Postmortem, both Coproscopy and Postmortem, and Immunological test), location of the study, season of data collection, and species of animals studied (Table 2) (Figure 3). Significant variation was observed across region of the study conducted, season, publication year, and animal species. The prevalence of ovine and caprine fasciolosis varied between different regions, ranging from 13.62% to 43.99%. The Amhara region subgroup possessed the highest prevalence (43.99%), while SNNPR region had the lowest prevalence (13.62%). The subgroup analysis based on sampling season showed that the highest infection rates were observed during Kiremt (equivalent to Winter) from June to August and Tsedey (equivalent to spring) from September to November (52.6%; 95% CI: 46.8–58.4%), compared to the lowest rates in Bega (equivalent to summer) from December to February (25.0%; 95% CI: 14.2–34.9%) (Table 2).

**Table 2.**
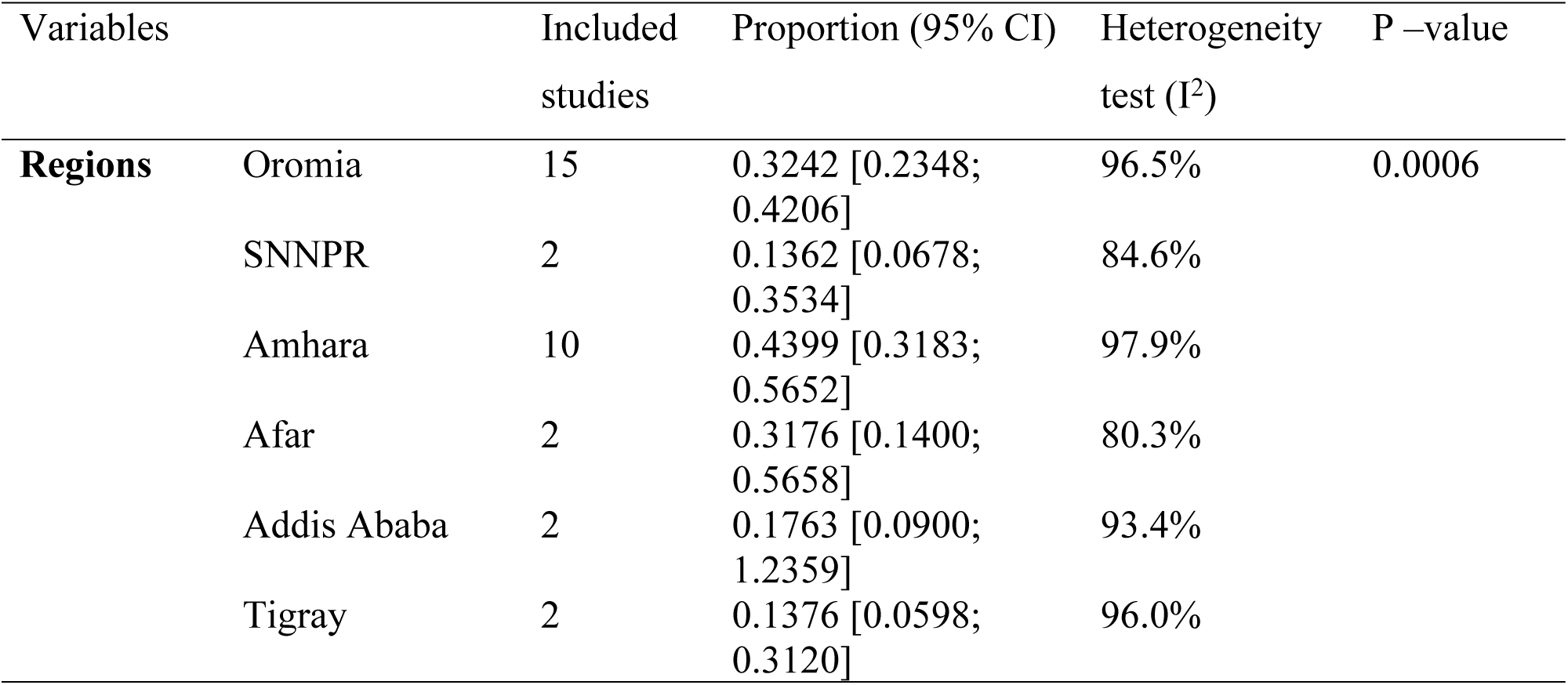
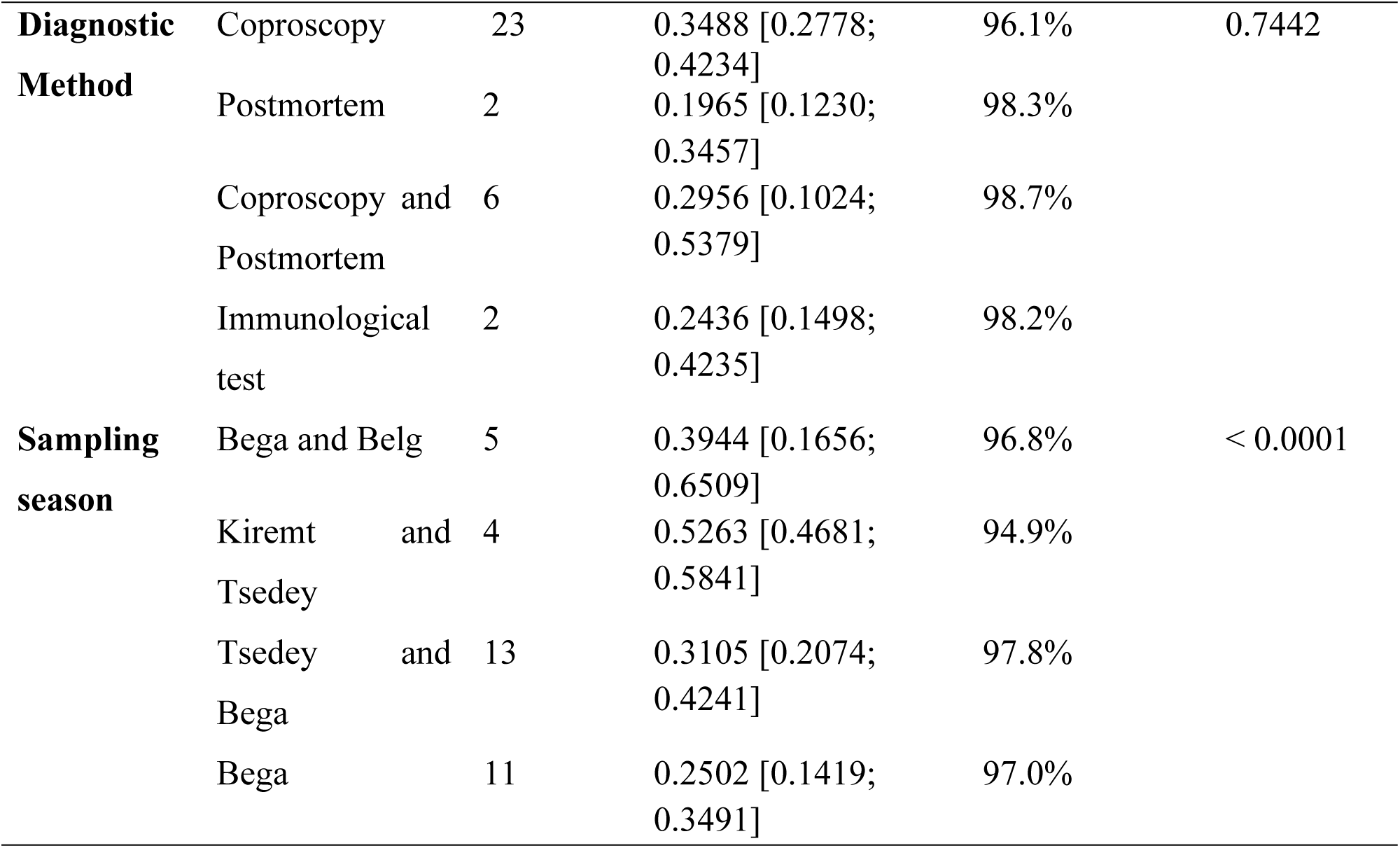
Subgroup analysis for diagnostic method, region, sampling season, and species of animals studied for fasciolosis.

The subgroup analysis based on species of animal showed that the disease prevalence in ovine was higher (37.18%; 95% CI: [31.06; 43.51]) than in caprine (12.76%; 95% CI: [4.06; 25.19]) (Figure 3).

**Figure 3:**
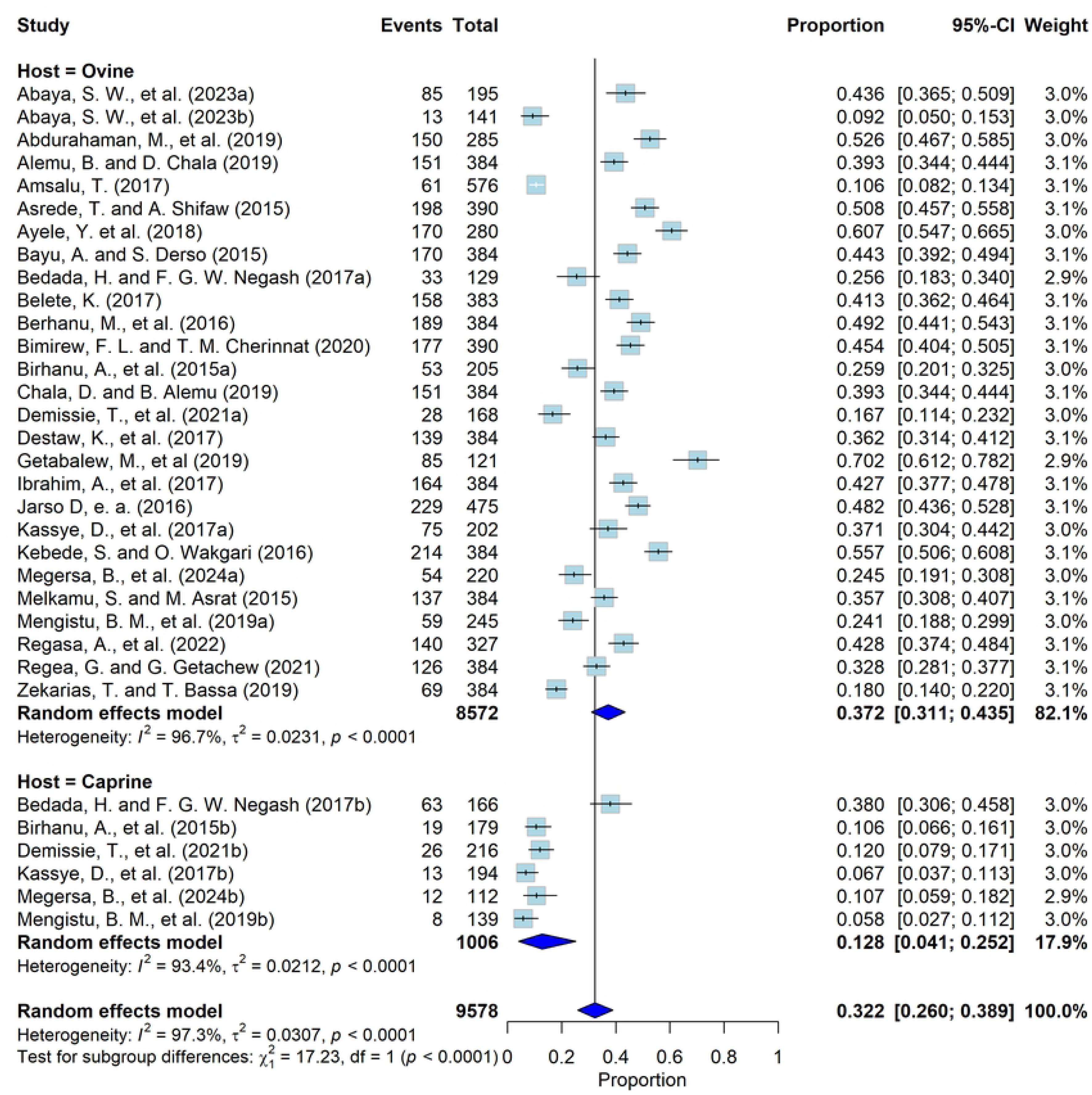
Subgroup analysis based on species of animal

#### 3.4.2. Sensitivity and influence analysis

Baujat diagnostic plots were used to detect studies that overly contribute to the heterogeneity in a meta-analysis. The plot shows the contribution of each study to the overall heterogeneity (as measured by Cochran’s Q) on the horizontal axis and its influence on the pooled effect size on the vertical axis[60]. Accordingly, a study by Amsalu, T. (2017) [34] contributed heavily to the overall heterogeneity in our meta-analysis (Figure 4)

**Figure 4:**
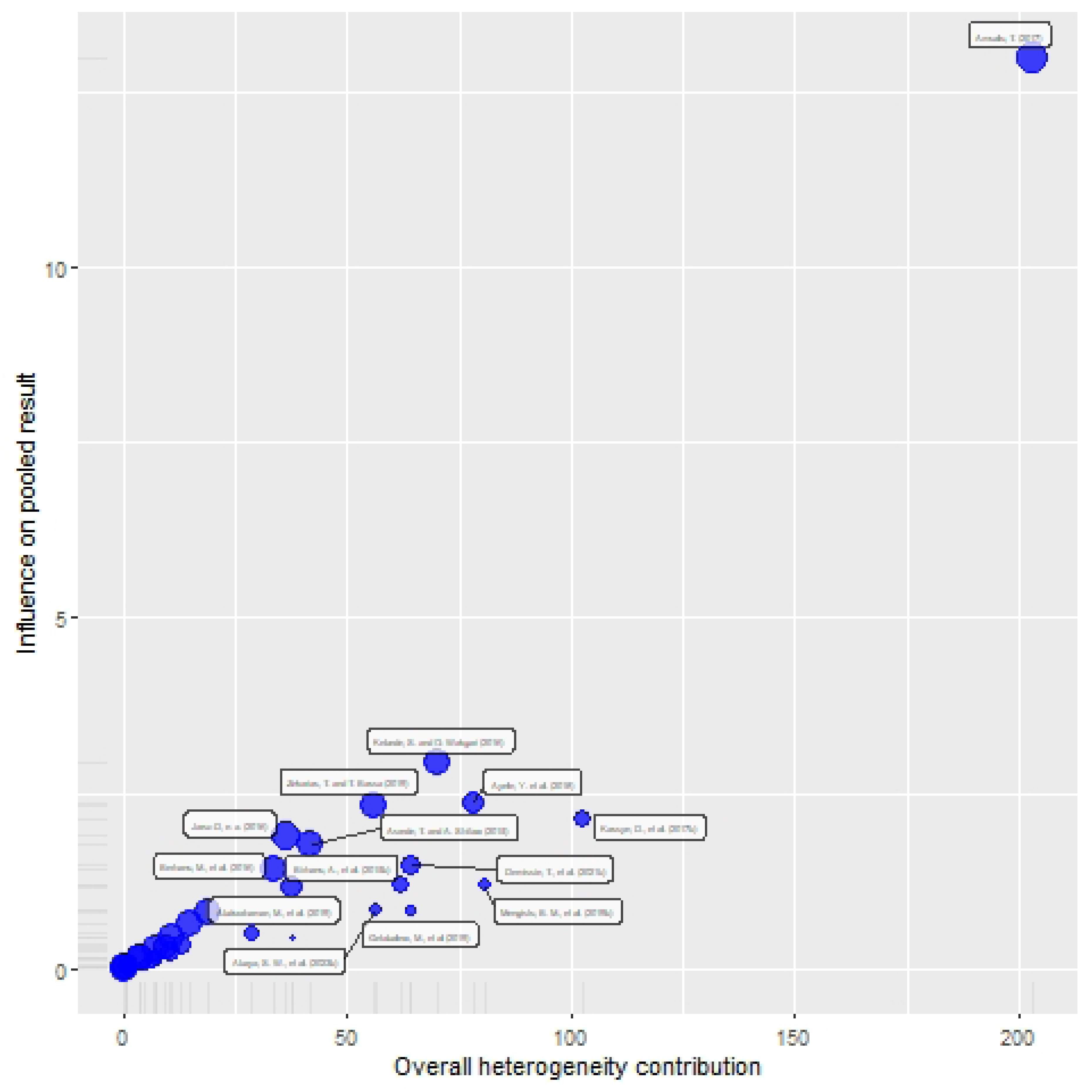
Baujat diagnostic plots of Fasciolosis prevalence among small ruminants in Ethiopia

A leave-one-out analysis was also performed to assess how each individual study influenced the overall effect size estimate. This involved systematically removing one study at a time and conducting a meta-analysis on the remaining (n-1) studies. If the confidence interval of the excluded study did not include the overall effect size estimate, it was considered to have a significant impact on the results [61]. The meta-analysis produced a pooled effect size of 0.322, which aligned with the confidence intervals of every individual study included. Sensitivity testing further confirmed the stability of these findings, as removing any single study from the analysis did not meaningfully change the overall prevalence estimate (Figure 5). Thus, the meta-analysis results encompassing all included studies in this study were reliable.

**Figure 5:**
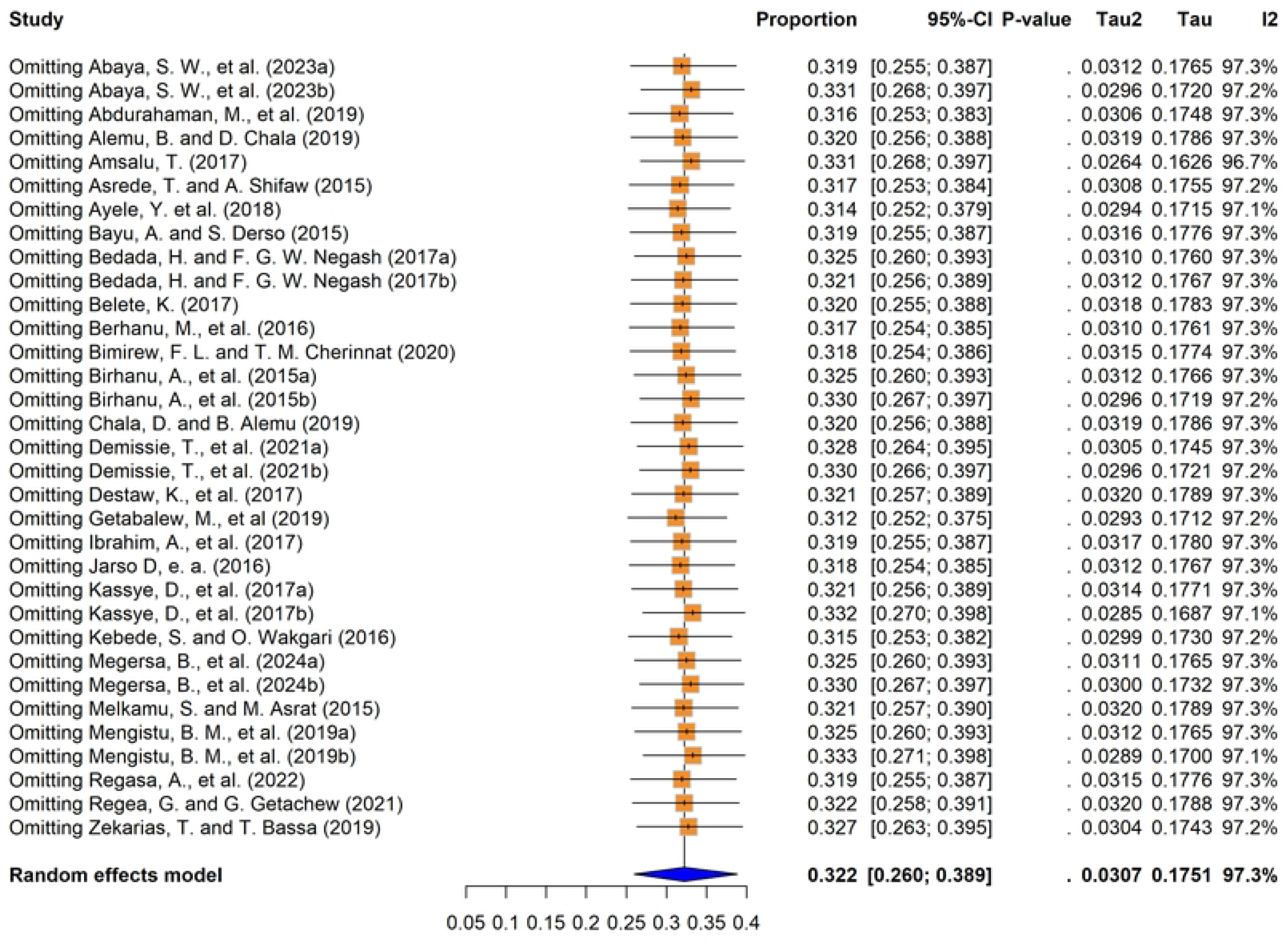
Sensitivity analysis of fasciolosis prevalence among small ruminants in Ethiopia

#### 3.4.3. Meta-regression models

Univariable and multivariable meta-regression models were employed to investigate sources of data heterogeneity. In univariable analysis, publication year and sample size were treated as continuous predictors, while study region, species, season, altitude, and diagnostic methods were analyzed as categorical variables within a mixed-effects framework. These predictors were evaluated for linear associations with effect sizes (the dependent variable). Animal species, geographic region, publication year, and sample size demonstrated statistically significant associations with variability across studies. Notably, animal species was found as the strongest explanatory factor, accounting for 25.17% of total heterogeneity (R² = 25.17%). The multivariable model further resolved 39.53% of heterogeneity. The Regression coefficients for multivariable model revealed that studies focusing on Ovine species (β = 0.2913) were associated with elevated effect sizes relative to Caprine studies after adjusting for covariates (Table 3).

**Table 3:**
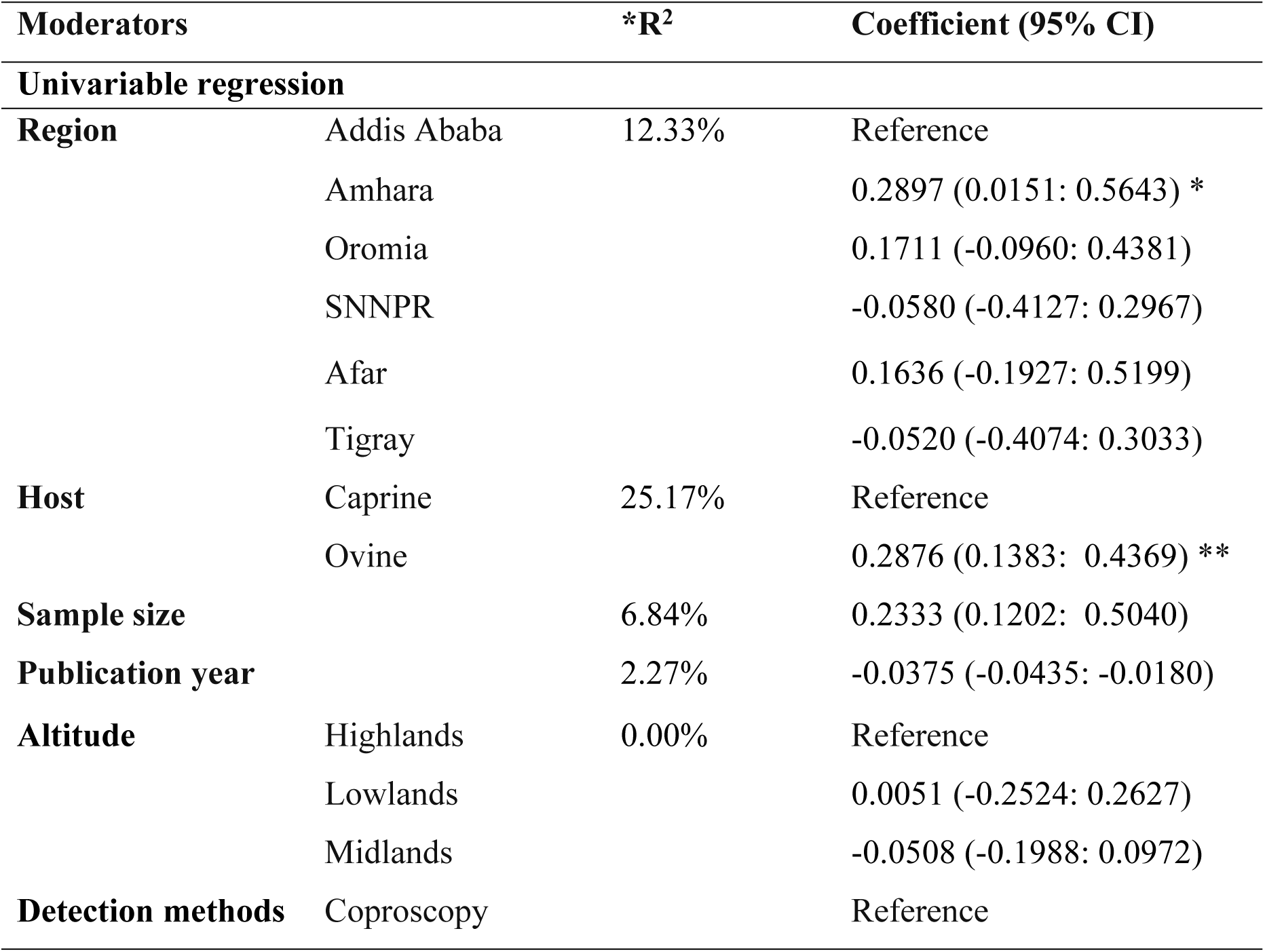
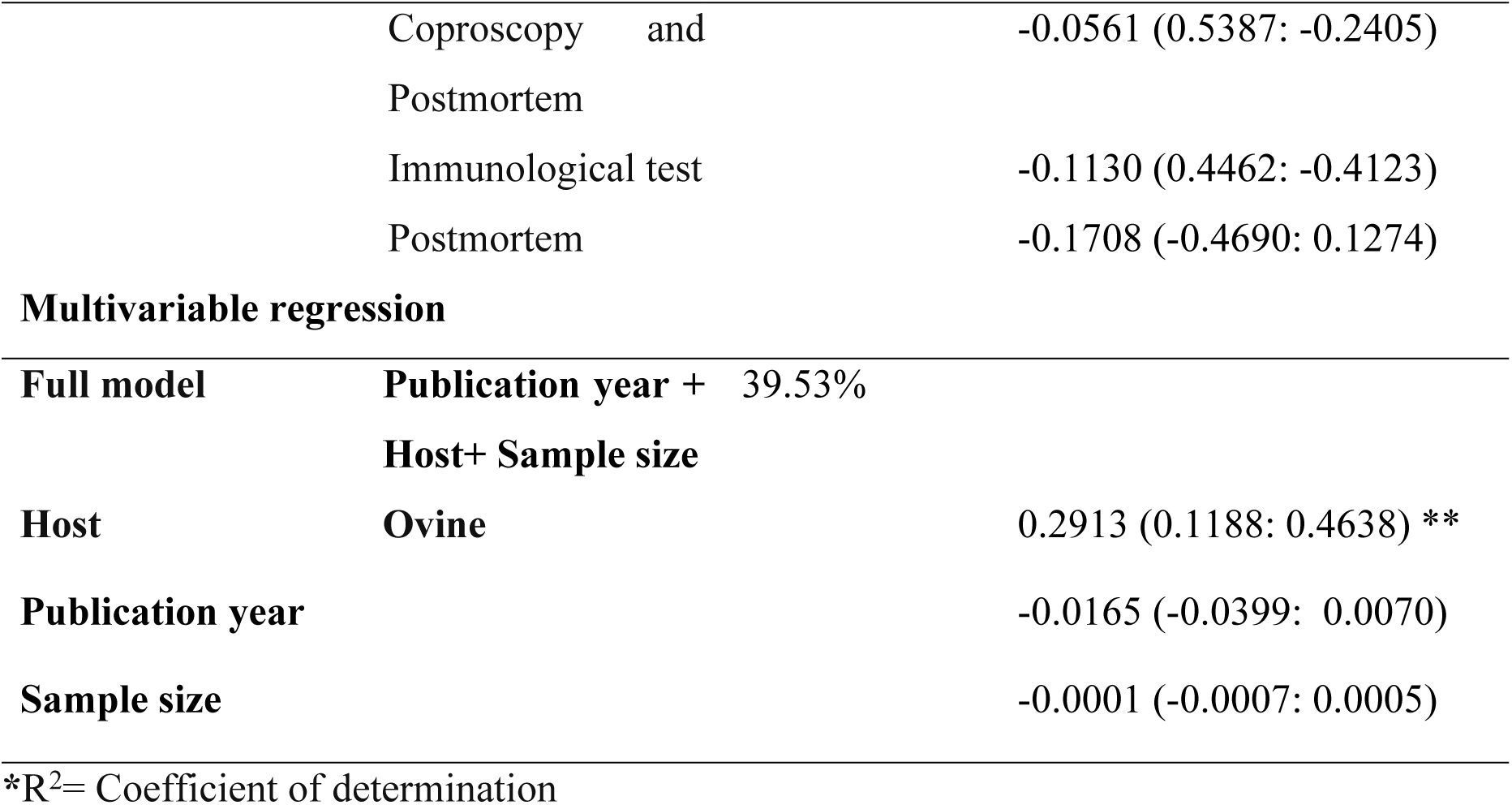
Uni-variable and multivariable meta-regression analysis results for the prevalence of fasciolosis among small ruminants in Ethiopia.

### 3.5. Publication bias

Publication bias was evaluated through visual and statistical methods [62]. Visual inspection of the funnel plot revealed a symmetrical distribution of study estimates (Figure 6). To complement this subjective assessment, quantitative statistical tests [63] were applied. Egger’s regression analysis (coefficient *t* = −1.75; p = 0.0906) and Begg’s rank correlation test (*z* = −2.17; p = 0.0599) both indicated no statistically significant asymmetry in the pooled data. These findings suggest no substantial evidence of publication bias or small-study effects influencing the meta-analytic results.

**Figure 6:**
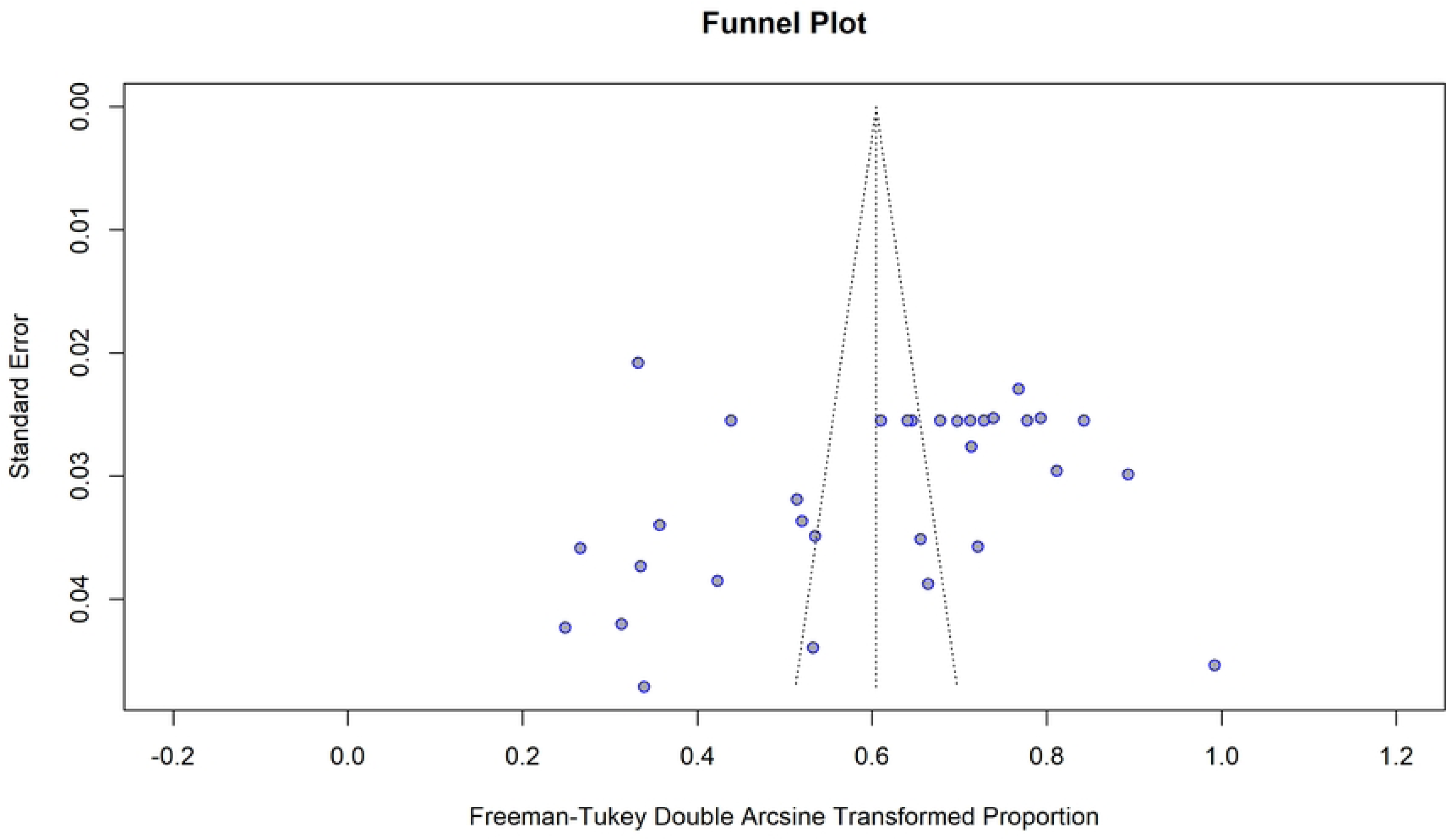
Funnel plot of the pooled prevalence of fasciolosis among small ruminants in Ethiopia

## 4. Discussion

Fasciolosis, a parasitic infection caused primarily by *Fasciola hepatica* and *Fasciola gigantica*, remains a significant concern in Ethiopia due to its widespread prevalence. In developing nations like Ethiopia, small ruminants, particularly sheep and goats serve as economically vital livestock, supporting livelihoods and food security. This emphasizes the need to synthesize existing data on fasciolosis prevalence and associated risk factors in these animals. To our knowledge, this study represents the first comprehensive systematic review and meta-analysis addressing the burden of fasciolosis in Ethiopian small ruminants, alongside an evaluation of contributing epidemiological variables. By consolidating current evidence, this work aims to support evidence-based interventions and inform targeted control strategies to mitigate the disease’s impact on livestock productivity and rural economies.

The findings of this study showed that the pooled prevalence of fasciolosis among small ruminants in Ethiopia was 32.25% (95% CI: 25.97–38.86). This result is larger than earlier studies conducted in China, which reported a prevalence of 26% [64], and a study conducted in Bangladesh, which reported a rate of 20% [65]. The higher prevalence of fasciolosis in Ethiopian small ruminants compared to other regions can be attributed to favorable ecological conditions for intermediate host snails [4], traditional management and husbandry practices [66], limited veterinary interventions and drug resistance [67], and differences in diagnostic methodologies that may affect reported prevalence rates [68].

Our analysis demonstrates significant geographic and species-specific variations in fasciolosis prevalence. The result of the region subgroup analysis showed that the highest prevalence of ovine and caprine fasciolosis was 43.99% [95% CI: 31.83; 56.52%] in Amhara region, and the lowest prevalence was observed in SNNPR (13.62% [95% CI: 6.78; 35.34%]). The significant variation in fasciolosis prevalence observed across regions and species may be explained by ecological, climatic, and husbandry-related factors that influence the transmission dynamics of the parasite. The Amhara region, characterized by high-altitude wetlands, perennial water bodies, and intensive irrigation systems (e.g., Lake Tana basin), provides optimal habitats for the intermediate hosts of *Fasciola* species [52].

The results indicate that sheep had a significantly higher prevalence of fasciolosis (37.18%; 95% CI: [31.06, 43.51]) compared to goats (12.76%; 95% CI: [4.06, 25.19])), suggesting species-specific susceptibility to the disease. This disparity may stem from differences in grazing behavior and physiological susceptibility. Sheep are known to graze closer to the ground and in wetter, marshy areas where *Fasciola* spp. larvae are more prevalent, increasing their exposure to infective metacercariae [69, 70]. In contrast, goats exhibit more selective browsing behavior, often feeding on shrubs and higher vegetation, which may reduce their risk of infection [71]. Additionally, goats may exhibit greater resistance to *Fasciola* infections due to variations in immune response or liver enzyme activity, as suggested by Molina-Hernández et al. (2015) [72]. These findings highlight the need for targeted control measures, particularly in sheep, to mitigate the impact of fasciolosis.

Seasonal subgroup analysis showed statistically significant variations in fasciolosis prevalence among small ruminants (p < 0.05), with the highest infection rates observed during Kiremt from June to August and Tsedey from September to November (52.6%; 95% CI: 46.8–58.4%), compared to the lowest rates in Bega from December to February (25.0%; 95% CI: 14.2–34.9%) (Table 2). This could be attributed to Ethiopia’s unique climatic and ecological drivers. Infection rates peaked during *Kiremt* (June–August) and *Tsedey* (September–November), coinciding with periods of elevated rainfall and humidity. These conditions promote the proliferation of the intermediate hosts for *Fasciola* species by creating saturated grazing areas and standing water bodies that serve as ideal habitats [73]. Additionally, moisture retention during *Tsedey* sustains vegetation contamination by metacercariae, prolonging livestock exposure [4]. In contrast, the arid *Bega* season (December– February) reduces transmission risk through habitat desiccation and cooler highland temperatures, which suppress snail activity and metacercarial survival. Shifts in livestock management, such as restricted grazing and stall-feeding during *Bega*, further limit contact with infected pastures [74].

However, this systematic review and meta-analysis has some limitations. First, the geographic scope was restricted to six of Ethiopia’s 11 administrative regions (Amhara, Oromia, SNNP, Tigray, Afar and Addis Ababa) due to lack of published data in underrepresented regions. This exclusion risks underrepresenting national epidemiological diversity, as ecological conditions (e.g., humidity, altitude) and livestock management practices in excluded regions may differ significantly, potentially biasing pooled prevalence estimates. Second, the time period inclusion criteria prioritized studies published after 2015 to align with recent epidemiological trends, but this approach may overlook historical data critical for contextualizing shifts in fasciolosis burden, such as evolving climate patterns, or changes in agricultural practices over decades. Finally, sparse data within subgroups such as regions, and climatic zones, likely diminished the statistical power and precision of subgroup analyses, resulting in broader confidence intervals and less definitive conclusions about niche epidemiological drivers. Collectively, these limitations highlight the challenges of synthesizing fragmented and heterogeneously reported data in Ethiopia, emphasizing the need for standardized, nationwide surveillance to improve future meta-analytic robustness.

## 5. Conclusions

This systematic review and meta-analysis emphasize fasciolosis as a pervasive threat to Ethiopian small ruminants, with a pooled national prevalence of 32.25%. The elevated burden in Ethiopia reflects the confluence of ecological, climatic, and management factors, including optimal conditions for *Lymnaea* snail proliferation, traditional grazing practices, and limited veterinary interventions. Policymakers, veterinarians, and researchers should prioritize evidence-based control programs targeting high-burden regions and seasonal variations through region-specific deworming, seasonal pasture management, and improved veterinary access for disease treatment, while expanding surveillance in underrepresented areas, conducting breed-specific resistance studies, and standardizing national reporting to address data gaps and climate-driven transmission risks.

## Abbreviations

CoCoPop: Condition, context, and population
*F. gigantica*: *Fasciola gigantica*
*F. hepatica*: *Fasciola hepatica*
PRISMA: Preferred Reporting Items for Systematic Review and Meta-Analysis
SNNPR: Southern Nations, Nationalities, and Peoples’ Region
WHO: World Health Organization

## Informed Consent Statement

We used published articles for further analysis. The participants of this study were articles. Consequently, consent was not obtained from the study participants.

## Ethics approval

Not applicable

## Data Availability Statement

All datasets are included in the manuscript or as supplementary files.

## Author contributions

Conceptualization: Simachew Getaneh Endalamew; Data curation: Simachew Getaneh Endalamew, Alebachew Tilahun Wassie, Andnet Yirga Assefa, Solomon Keflie Assefa, Solomon Mekuriaw Ayalew, Yihenew Getahun Ambaw; Software, Methodology and formal analysis: Simachew Getaneh Endalamew, Solomon Keflie Assefa; Supervision and Validation: Alebachew Tilahun Wassie, Solomon Keflie Assefa, Yihenew Getahun Ambaw, Solomon Mekuriaw Ayalew, Andnet Yirga Assefa; Writing-original draft: Simachew Getaneh Endalamew; Writing-review & editing: Simachew Getaneh Endalamew, Alebachew Tilahun Wassie, Solomon Keflie Assefa, Yihenew Getahun Ambaw, Solomon Mekuriaw Ayalew, Andnet Yirga Assefa

## Acknowledgements

We thank Bahir Dar university, School of Veterinary Medicine for the free internet access to facilitate this research work.

## Reference

1. Tsega, M., S. Dereso, and A. Getu, A review on ruminant fasciolosis. Open Access Library Journal, 2015. 2(8): p. 1–8.

2. Dalton, J.P., Fasciolosis. 2021: CABI.

3. Mas-Coma, S., M.A. Valero, and M.D. Bargues, Human and animal fascioliasis: origins and worldwide evolving scenario. Clinical microbiology reviews, 2022. 35(4): p. e00088–19.

4. Yilma, J. and J. Malone, A geographic information system forecast model for strategic control of fasciolosis in Ethiopia. Veterinary parasitology, 1998. 78(2): p. 103–127.

5. Tolan Jr, R.W., Fascioliasis due to Fasciola hepatica and Fasciola gigantica infection: an update on this ‘neglected’neglected tropical disease. Laboratory Medicine, 2011. 42(2): p. 107–116.

6. Siles-Lucas, M., D. Becerro-Recio, J. Serrat, and J. González-Miguel, Fascioliasis and fasciolopsiasis: Current knowledge and future trends. Research in veterinary science, 2021. 134: p. 27–35.

7. Mas-Coma, S., M.A. Valero, and M.D. Bargues, Fasciola and Fasciolopsis. Biology of Foodborne Parasites, 2015: p. 371–404.

8. World Health Orginizatio (WHO). Neglected Tropical Diseases. 2023 Available from: https://www.who.int/health-topics/neglected-tropical-diseases.

9. Flores-Velázquez, L.M., M.T. Ruiz-Campillo, G. Herrera-Torres, Á. Martínez-Moreno, F.J. Martínez-Moreno, R. Zafra, L. Buffoni, P.J. Rufino-Moya, V. Molina-Hernández, and J. Pérez, Fasciolosis: pathogenesis, host-parasite interactions, and implication in vaccine development. Frontiers in veterinary science, 2023. 10: p. 1270064.

10. Alba, A., A.A. Vazquez, and S. Hurtrez-Boussès, Towards the comprehension of fasciolosis (re-) emergence: an integrative overview. Parasitology, 2021. 148(4): p. 385–407.

11. Köstenberger, K., A. Tichy, K. Bauer, P. Pless, and T. Wittek, Associations between fasciolosis and milk production, and the impact of anthelmintic treatment in dairy herds. Parasitology research, 2017. 116: p. 1981–1987.

12. Mehmood, K., H. Zhang, A.J. Sabir, R.Z. Abbas, M. Ijaz, A.Z. Durrani, M.H. Saleem, M.U. Rehman, M.K. Iqbal, and Y. Wang, A review on epidemiology, global prevalence and economical losses of fasciolosis in ruminants. Microbial pathogenesis, 2017. 109: p. 253–262.

13. Fennouh, C., M. Nabi, I. Ouchetati, O. Salhi, N. Ouchene, H. Dahmani, A. Haif, D. Mokrani, and N.K. Touhami, A comprehensive analysis of fasciolosis prevalence and risk factors in humans and animals: First report in Algeria. Journal of Helminthology, 2025. 99: p. e26.

14. Arbabi, M., E. Nezami, H. Hooshyar, and M. Delavari, Epidemiology and economic loss of fasciolosis and dicrocoeliosis in Arak, Iran. Veterinary world, 2018. 11(12): p. 1648.

15. Cwiklinski, K., S. O’neill, S. Donnelly, and J. Dalton, A prospective view of animal and human Fasciolosis. Parasite immunology, 2016. 38(9): p. 558–568.

16. Beesley, N., C. Caminade, J. Charlier, R. Flynn, J. Hodgkinson, A. Martinez-Moreno, M. Martinez-Valladares, J. Perez, L. Rinaldi, and D. Williams, Fasciola and fasciolosis in ruminants in Europe: Identifying research needs. Transboundary and emerging diseases, 2018. 65: p. 199–216.

17. Tolosa, T. and W. Tigre, The prevalence and economic significance of bovine fasciolosis at Jimma abattoir, Ethiopia. The Internet Journal of Veterinary Medicine, 2007. 3(2): p. 1–7.

18. Ashrafi, K., M. Valero, M. Panova, M. Periago, J. Massoud, and S. Mas-Coma, Phenotypic analysis of adults of Fasciola hepatica, Fasciola gigantica and intermediate forms from the endemic region of Gilan, Iran. Parasitology International, 2006. 55(4): p. 249–260.

19. Zerna, G., T.W. Spithill, and T. Beddoe, Current status for controlling the overlooked caprine fasciolosis. Animals, 2021. 11(6): p. 1819.

20. Munn, Z., S. Moola, K. Lisy, D. Riitano, and C. Tufanaru, Methodological guidance for systematic reviews of observational epidemiological studies reporting prevalence and cumulative incidence data. JBI Evidence Implementation, 2015. 13(3): p. 147–153.

21. Parums, D.V., Review articles, systematic reviews, meta-analysis, and the updated preferred reporting items for systematic reviews and meta-analyses (PRISMA) 2020 guidelines. Medical science monitor: international medical journal of experimental and clinical research, 2021. 27: p. e934475–1.

22. Harrer, M., P. Cuijpers, T. Furukawa, and D. Ebert, Doing meta-analysis with R: A hands-on guide. 2021: Chapman and Hall/CRC.

23. DerSimonian, R. and N. Laird, Meta-analysis in clinical trials revisited. Contemporary clinical trials, 2015. 45: p. 139–145.

24. Doi, S.A., J.J. Barendregt, S. Khan, L. Thalib, and G.M. Williams, Advances in the meta-analysis of heterogeneous clinical trials I: the inverse variance heterogeneity model. Contemporary clinical trials, 2015. 45: p. 130–138.

25. IntHout, J., J.P. Ioannidis, and G.F. Borm, The Hartung-Knapp-Sidik-Jonkman method for random effects meta-analysis is straightforward and considerably outperforms the standard DerSimonian-Laird method. BMC medical research methodology, 2014. 14: p. 1–12.

26. Röver, C., G. Knapp, and T. Friede, Hartung-Knapp-Sidik-Jonkman approach and its modification for random-effects meta-analysis with few studies. BMC medical research methodology, 2015. 15: p. 1–7.

27. Jackson, D. and J. Bowden, Confidence intervals for the between-study variance in random-effects meta-analysis using generalised heterogeneity statistics: should we use unequal tails? BMC medical research methodology, 2016. 16: p. 1–15.

28. Chochran, W., Some methods for strengthening the common chi-squared tests. Biometrics, 1954. 10(4): p. 417–451.

29. Higgins, J.P. and S.G. Thompson, Quantifying heterogeneity in a meta-analysis. Statistics in medicine, 2002. 21(11): p. 1539–1558.

30. IntHout, J., J.P. Ioannidis, M.M. Rovers, and J.J. Goeman, Plea for routinely presenting prediction intervals in meta-analysis. BMJ open, 2016. 6(7): p. e010247.

31. Abaya, S.W., S.T. Mereta, F.D. Tulu, Z. Mekonnen, M. Ayana, M. Girma, H.R. Vineer, S.M. Mor, C. Caminade, and J. Graham-Brown, Prevalence of human and animal fasciolosis in Butajira and Gilgel Gibe health demographic surveillance system sites in Ethiopia. Tropical Medicine and Infectious Disease, 2023. 8(4): p. 208.

32. Abdurahaman, M., T. Dinagde, T. Kedir, T. Ahimad, T. Said, T. Mamo, and H. Tesfaye, A Study on Prevalence of Ovine Fasciolosis in Busa Town, Dawo Woreda, South West Shoa Zone, Oromia Region. International Journal of Research, 2019. 7(3): p. 1–6.

33. Alemu, B. and D. Chala, Prevalence of ovine fasciolosis and its economic impacts in and around Ambo, Ethiopia. International Journal, 2019. 5(1): p. 033–038.

34. Amsalu, T., Prevalence of ovine fasciolosis and loss due to liver condemnation at Bahir Dar Town, Ethiopia. Adv. Biol. Res, 2017. 11(5): p. 286–294.

35. Asrede, T. and A. Shifaw, Coprological Study on the prevalence of Ovine Fasciolosis in Debre Birhan Agricultural Research Center, Ethiopia. Eur J Biol Sci, 2015. 7: p. 103–7.

36. Ayele Y., w.F., and Yeshiwas T, The Prevalence of Bovine and Ovine Fasciolosis and the Associated Economic Loss Due to Liver Condemnation in and around Debire Birhan, Ethiopia. . SOJ Immunol 2018. 6 (3): 1–11.

37. Bayu, A. and S. Derso, Prevalence of Ovine Fasciolosis and its Associated Risk Factorin and Around Debre Elias District, East Gojam, North West of Ethiopia. 2015.

38. Bedada, H., F. Gizaw, W. Negash, A. Hadush, A. Wassie, and A. Gebregergious, Epidemiology of Small Ruminant Fasciolosis in Arid Areas of Lower Awash River Basin, Afar Region, Ethiopia. Animal and Veterinary Sciences, 2017. 5(6): p. 102–107.

39. Belete, K., A cross sectional study on the coprological prevalence of ovine fasciolosis in Amhara Sayint District, Ethiopia. Journal of Veterinary Medicine Research, 2017. 4(6): p. 1092.

40. Berhanu, M., G. Ayalew, and A. Tilahun, Prevalence and Associated Risk Factors for Ovine Fasciolosis in Selected Areas of North Gondar, Ethiopia. Advances in Biological Research, 2016. 10(3): p. 162–166.

41. Bimirew, F.L. and T.M. Cherinnat, Coprological Examination of Ovine Fasciolosis in Horro District Community Based Sheep Breeding Program, Horro Guduru Wollega Zone, Western Ethiopia. Journal of Veterinary Healthcare, 2020. 2(2): p. 31–38.

42. Birhanu, A., R. Tesfaye, and S. Derso, Prevalence and associated risk factors of Fasciola infection in small ruminants slaughtered at Addis Ababa abattoir Enterprise, Ethiopia with reference to diagnostic value of its coprological examination. African Journal of Basic & Applied Sciences, 2015. 7(4): p. 181–186.

43. Chala, D. and B. Alemu, Prevalence of Ovine Fasciolosis and Its Economic Loss In and Around Ambo, Ethiopia. 2019.

44. Dabasa, G., T. Shanko, W. Zewdei, K. Jilo, G. Gurmesa, and N. Abdela, Prevalence of small ruminant gastrointestinal parasites infections and associated risk factors in selected districts of Bale zone, south eastern Ethiopia. Journal of Parasitology and Vector Biology, 2017. 9(6): p. 81–88.

45. Demissie, T., B. Wakjira, J. Shiferaw, A. Feyisa, and Y.H. Tolossa, Pre-slaughter coproscopic and abattoir prevalence of GIT parasites of sheep and goats at Bishoftu ELFORA export abattoir, Oromia, Ethiopia.

46. Destaw, K., W. Mitku, M. Hamid, B. Alemu, and T. Tintagu, Prevalence of ovine fasciolosis in selected Kebeles of Wogera District, North Gondar zone, Ethiopia. Int. J. Adv. Res. Biol. Sci, 2017. 4(8): p. 78–84.

47. Getabalew, M., T. Alemneh, D. Akeberegn, A. Getie, B. Fekadu, H. Hadgu, and N. Zeselase, Prevalence of Ovine Fasciolosis in Debre Berhan Agricultural Research Center, North Shewa Zone, Ethiopia.

48. Ibrahim, A., D. Nolkes, E. Gezahegn, and M. Taye, Prevalence of ovine fasciolosis in Jimma and selected rural kebeles near Jimma, Southwest Ethiopia. J Vet Sci Technol, 2017. 8(424): p. 2.

49. Jarso D, e.a., Study on Prevalence of Ovine Fasciolosis in and Around Debre Berhan Sheep Breeding and Forage Multiplication Center. Journal of Veterinary Science & Research, 2016.

50. Kassye, D., M. Gebeyehu, and D. Mekonnen, Prevalence and associated risk factors of small ruminant Fasciolosis in Haramaya district, Eastern Ethiopia. Acta Parasitol, 2017. 8: p. 144–149.

51. Kebede, S. and O. Wakgari, The prevalence study of ovine fasciolosis in Jima Rare District, Horo Guduru Wollega Zone, Oromia Regional State, Western Ethiopia. 2016.

52. Megersa, B., B. Hussien, J. Shemsu, R. Kassahun, O. Merera, N. Moje, B.M. Edao, H. Ashenafi, and D. Ayana, Trematode Infestations in Ruminants and Their Snail Hosts across varied Agro-Ecological zones in Ethiopia: Implication for Public Health Risk. 2024.

53. Melkamu, S. and M. Asrat, Study on the prevalence of Ovine fasciolosis in Ambasel Woreda, South Wollo zone, Amhara regional state, Ethiopia. Journal of Animal Research, 2015. 5(3): p. 437–441.

54. Mengistu, B.M., A.A. Azbite, and H.K. Bitsue, Prevalence of Small Ruminants Fasciolosis in Mekelle, Tigrai Regional State, Northern Ethiopia. Journal of Agriculture and Ecology Research International, 2019. 20(1): p. 1–8.

55. Regasa, A., B. Mesfin, and M. Adem, Study on prevalence and risk factors of Ovine Fasciolosis on Shirka District, Arsi Zone, Eastern Ethiopia. J Vet Med Heal, 2022. 6: p. 2.

56. Regea, G. and G. Getachew, Study on prevalence and associated risk factors of ovine fasciolosis in and around Nekemte Town, Oromia, Ethiopia. Journal of Veterinary Medicine and Animal Sciences, 2021. 4(1).

57. Zekarias, T. and T. Bassa, Prevalence of Ovine Fasciolosis in Damot Sore Woreda, Wolayta Zone, Ethiopia. 2019.

58. Spineli, L.M. and N. Pandis, Prediction interval in random-effects meta-analysis. American journal of orthodontics and dentofacial orthopedics, 2020. 157(4): p. 586–588.

59. Ades, A., G. Lu, and J. Higgins, The interpretation of random-effects meta-analysis in decision models. Medical Decision Making, 2005. 25(6): p. 646–654.

60. Baujat, B., C. Mahé, J.P. Pignon, and C. Hill, A graphical method for exploring heterogeneity in meta-analyses: application to a meta-analysis of 65 trials. Statistics in medicine, 2002. 21(18): p. 2641–2652.

61. Hedges, L.V. and I. Olkin, Statistical methods for meta-analysis. 2014: Academic press.

62. Schwarzer, G., J.R. Carpenter, G. Rücker, G. Schwarzer, J.R. Carpenter, and G. Rücker, Small-study effects in meta-analysis. Meta-analysis with R, 2015: p. 107–141.

63. Egger, M., G.D. Smith, M. Schneider, and C. Minder, Bias in meta-analysis detected by a simple, graphical test. Bmj, 1997. 315(7109): p. 629–634.

64. Lan, Z., J. Yu, X. Zhang, A. Zhang, R. Deng, B. Li, Q. Lv, X. Ma, J. Gao, and C. Wang, Prevalence and risk factors of ovine and caprine fasciolosis in the last 20 years in China: A systematic review and meta-analysis. Animals, 2023. 13(10): p. 1687.

65. Mia, M.M., M. Hasan, and M.R. Chowdhury, A systematic review and meta-analysis on prevalence and epidemiological risk factors of zoonotic fascioliasis infection among the ruminants in Bangladesh. Heliyon, 2021. 7(12).

66. Dargie, J., The impact on production and mechanisms of pathogenesis of trematode infections in cattle and sheep. International journal for parasitology, 1987. 17(2): p. 453–463.

67. Howell, A., M. Baylis, R. Smith, G. Pinchbeck, and D. Williams, Epidemiology and impact of Fasciola hepatica exposure in high-yielding dairy herds. Preventive veterinary medicine, 2015. 121(1-2): p. 41–48.

68. Mezo, M., M. González-Warleta, J.A. Castro-Hermida, and F.M. Ubeira, Evaluation of the flukicide treatment policy for dairy cattle in Galicia (NW Spain). Veterinary parasitology, 2008. 157(3-4): p. 235–243.

69. Foreyt, W.J., Diagnostic parasitology. Veterinary Clinics of North America: Small Animal Practice, 1989. 19(5): p. 979–1000.

70. Torgerson, P. and J. Claxton, Epidemiology and control. Fasciolosis, 1999. 113: p. 149.

71. Hansen, J. and B. Perry, The epidemiology, diagnosis and control of helminth parasites of ruminants. 1994.

72. Ruiz-Campillo, M.T., V. Molina Hernandez, A. Escamilla, M. Stevenson, J. Perez, A. Martinez-Moreno, S. Donnelly, J.P. Dalton, and K. Cwiklinski, Immune signatures of pathogenesis in the peritoneal compartment during early infection of sheep with Fasciola hepatica. Scientific reports, 2017. 7(1): p. 2782.

73. Sissay, M.M., A. Uggla, and P.J. Waller, Prevalence and seasonal incidence of nematode parasites and fluke infections of sheep and goats in eastern Ethiopia. Tropical Animal Health and Production, 2007. 39: p. 521–531.

74. Yilma, J. and A. Mesfin, Dry season bovine fasciolosis in Northwestern part of Ethiopia. Revue de médecine vétérinaire, 2000. 151(6): p. 493–500.

